# Building neuroanatomical resources for three-spined sticklebacks: Brain areas important for social behavior

**DOI:** 10.64898/2026.01.02.697386

**Authors:** Tina Barbasch, Usan Dan, Catherine Marquez, Julia Ciura, Meghan F. Maciejewski, Makoto Kusakabe, Naoyuki Yamamoto, Hiyu Kanbe, Jun Kitano, Alison M. Bell

**Affiliations:** School of Integrative Biology, University of Illinois Urbana Champaign, Urbana, IL, USA; Carl R. Woese Institute for Genomic Biology, University of Illinois Urbana Champaign, Urbana, IL, USA; Neuroscience Program, University of Illinois Urbana Champaign, Urbana, IL, USA; School of Molecular and Cellular Biology, University of Illinois Urbana Champaign, Urbana, IL, USA; Creative Science Unit, Faculty of Science, Shizuoka University, 836 Ohya, Suruga-ku, Shizuoka, 422-8529, Japan; Laboratory of Fish Biology, Graduate School of Bioagricultural Sciences, Nagoya University, Furo-cho, Chikusa-ku, Nagoya, 464-8601, Japan; Ecological Genetics Laboratory, National Institute of Genetics, Mishima, Shizuoka 411-8540, Japan

**Keywords:** Brain atlas, social decision-making network, three-spined stickleback (*Gasterosteus aculeatus*), dopaminergic system

## Abstract

**Introduction:** Three-spined stickleback fish are famous for their diversity and charismatic social behavior. However, there are few neuroanatomical resources for studying the neural and brain mechanisms underlying their fascinating behavior.

**Methods and Results:** We identify 11 brain areas important for social behavior by referencing brain atlases for six other teleost fishes. Brain regions were identified via neuroanatomical landmarks and we characterized the presence / absence of tyrosine hydroxylase (TH), a key gene product of the dopaminergic system, in those regions. Comparing the neuroanatomical location of these regions in the stickleback brain and the expression of TH therein to that of other fish species highlights similarities and differences and the need for a brain atlas specific to sticklebacks. This resource serves as a map of the location of regions important for social behavior in the stickleback brain.

**Conclusion:** This resource will help guide future studies connecting gene function to social behavior through the brain and will enable future work to understand the evolution of neural mechanisms that contribute to the diversity of social behavior in this emerging model organism.

## Introduction

Three-spined stickleback (*Gasterosteus aculeatus*) have been a favorite animal model for studies of animal behavior since the early ethologists, including Nobel laureate Niko Tinbergen [1–3]. Seminal studies in this organism on sign stimuli, fixed action patterns, displacement behavior and drive [4], for example, helped to define the emerging field of ethology in the early 20th century. Sticklebacks have remained popular subjects ever since, with hundreds of publications and multiple book volumes [3,5–8] devoted to them, and there is a vibrant worldwide community of researchers across dozens of labs studying the behavior of this small fish.

However, there are relatively few neuroanatomical tools and resources available for tackling the neural basis of the behavior that initially made sticklebacks so famous. For example, there is not yet a brain atlas available which delineates where regions are located throughout the stickleback brain. Brain atlases enable collaboration and ensure consistency across different labs by providing a reference that delineates and names different brain areas. A map of the stickleback brain will allow researchers to compare the diversity in brain structure across populations of sticklebacks and to other species, and will enable functional studies of genes, brain, and behavior by connecting gene function to behavior through the brain. Neuroanatomical resources specific to sticklebacks are needed because the unique outward folding of the teleost pallium during development – “eversion” – leads to an altered arrangement of pallial zones compared to the “evaginated” brains of other vertebrates, making homology difficult across vertebrates [9,10]. Although there are excellent anatomically defined brain atlases available for other small fishes, such as zebrafish [11–13], brain organization varies considerably among fishes [14,15], highlighting the need for neuroanatomical resources specific to stickleback.

Given their rich behavioral repertoire, distinguished ethological history, behavioral diversity within and among the populations, and the worldwide community of researchers studying them, sticklebacks have the potential to become an important model in evolutionary neuroscience. Here, we contribute to this effort by identifying brain areas important for social behavior in sticklebacks. We focus on brain areas important for social behavior because of the rich history and literature on social and reproductive behaviors such as aggression, parental care and courtship in sticklebacks [16] and characterize the hypothesized vertebrate social decision-making network (SDMN), a set of interconnected brain regions with conserved functions related to various social behaviors and reward processing [17–21] in vertebrates. This resource serves as an initial step toward developing an eventual brain-wide atlas for stickleback.

We compare cresyl-violet stained sections of stickleback brain to the brain atlases of other fish species including zebrafish [11], guppy [22] and cichlids [23,24] and use neuroanatomical landmarks to identify hypothesized SDMN nodes. Gene products of the dopaminergic system such as tyrosine hydroxylase (TH, the enzyme limiting dopamine synthesis) play a fundamental role in regulating complex social behavior in all vertebrate taxa and are a hallmark of hypothesized SDMN nodes [19], therefore we also assess evidence for tyrosine hydroxylase staining in the hypothesized brain areas. This atlas will serve as a resource for neuroscientific studies of social behavior in this emerging model organism.

## Methods

### Neuroanatomical SDMN delineations

Brain tissue for the atlas was obtained from anadromous three-spined stickleback (*G. aculeatus*) originating from a spawning site in Bekanbeushi River, Akkeshi, Hokkaido (see [25,26]). Two genetically divergent forms (‘Japan Sea’ and ‘Pacific Ocean’) are present in this population, thus it is important to note that the individual used to generate this atlas was of the Pacific Ocean form. Fish were housed in the laboratory in 10% seawater (Instant Ocean, Aquarium Systems, OH, USA) prior to tissue collection. A single adult male individual was euthanized by immersing in a high dosage of anesthesia (tricane methansulfonate, 300mg/L), followed by perfusion through the heart with 2% paraformaldehyde and 1% glutaraldhyde in 0.1M phosphate buffer (pH7.4). The dissected brain was post-fixed in a fresh solution of the same fixative overnight and was then immersed in 20% sucrose for cryoprotection. The brain was embedded in 5% agarose (Sigma, Type IX), frozen in -60ºC n-hexane, and cryosectioned at 30 μm transversely. Sections were mounted serially on gelatin-coated glass slides.

Nissl staining was performed using cresyl violet. Subsequent to washes in 0.1M phosphate buffered saline (PBS) to remove agar, the sections were dehydrated through an ascending series of ethanol and then rehydrated through a descending series of ethanol. Then, the sections were stained with 0.1% cresyl violet, dehydrated in an ascending series of ethanol, cleared in xylene, and coverslipped with Permount (Fisher Scientific, Hampton, NH, USA) for imaging. The sections were observed under a light microscope (BX60; Olympus), and photomicrographs were taken by a digital camera (DP-70; Olympus), which was controlled by cellSens software (Olympus).

We referenced published brain atlases for zebrafish (*Danio rerio*) [11], guppy (*Poecilia reticulata*) [22], and cichlid (*Astatotilapia burtoni*) [23,24] to aid in identifying brain regions. Brain regions were identified via neuroanatomical landmarks. We focused on 11 of the 12 regions within the hypothesized vertebrate social decision-making network (excluding the ventral pallidum as there is no known or putative teleost homolog) [19] (Table 1). To facilitate comparisons across teleost species and provide a more general resource for future study of the SDMN, we additionally referenced brain atlases for sea bass (*Dicentrarchus labrax*) [27–29], tilapia (*Oreochromis mossambicus*) [30], and cardinal tetra (*Paracheirodon axelrodi*) [31] in creating the table. For each species, we identified regions (and sub-regions, when delineated) that most closely corresponded to each of our delineations in stickleback, based on neuroanatomical similarity.

**Table 1.**
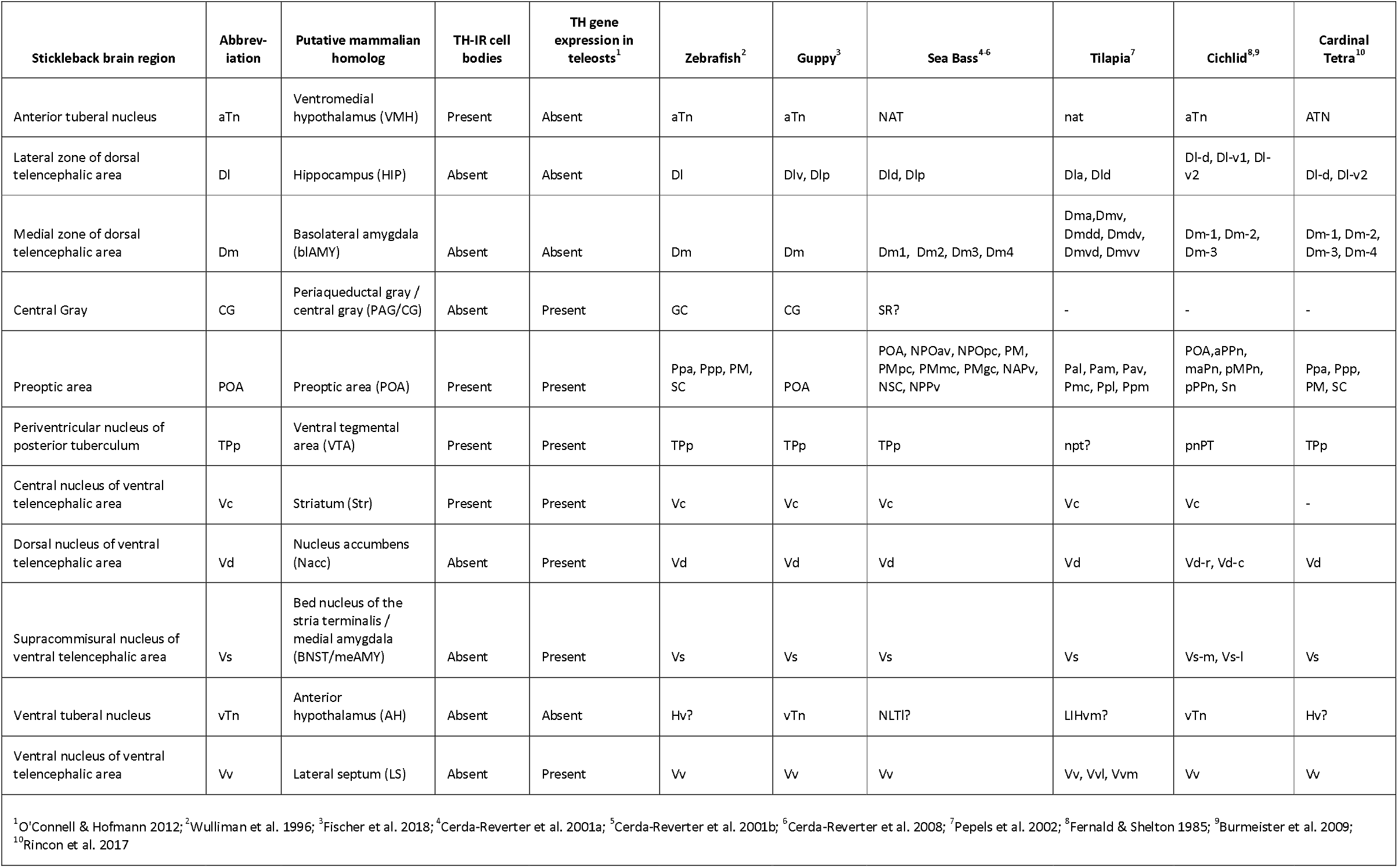
List of brain regions important for social behavior delineated in stickleback (this study) with putative mammalian homolog and presence / absence of TH staining in each region. Also listed are the corresponding region or set of regions in brain atlases of six other teleost fishes. Regions were identified that most closely align anatomically with each stickleback region delineated here, to highlight differences in nomenclature and build a resource to facilitate comparisons across species. Dashes represent regions where no clear corresponding region was found, based on similarity in anatomical location, and a question mark after a region represents places where a seemingly corresponding region was identified but nomenclature was for a different region, and thus the correspondence is uncertain.

### Tyrosine hydroxylase quantification

Tyrosine hydroxylase immunoreactive cell bodies were quantified in brain tissue collected from two lab-reared marine stickleback originating from New Brunswick, Canada or Canal Lake, Nova Scotia, Canada, and maintained at the University of Illinois Urbana Champaign. While there is likely to be subtle differences in brain connectivity and activity among behaviorally divergent stickleback populations, we do not expect overall brain architecture and organization to differ. Individuals used in this study were euthanized by rapid decapitation and whole brains (N = 2) were dissected and fixed in 4% paraformaldehyde at 4°C for five hours. Fixed brains were then rinsed in 1X PBS and dehydrated in 30% sucrose. After dehydration, brains were embedded in Tissue-Tek® O.C.T. Compound (Electron Microscopy Sciences, Hatfield, PA, USA) and frozen at -80°C until sectioning. Brains (approx. 3 × 3 × 7 mm in size) were cryosectioned at 20 μm along the coronal (transverse) plane and sections were placed serially on charged slides (Superfrost Plus, VWR, Randor, PA, USA). To compare the distribution of tyrosine hydroxylase throughout the SDMN to that of other teleosts, the presence of TH positive cells across all 11 brain regions was assessed using a standard fluorescence immunohistochemistry protocol. Briefly, slides were incubated for 1 hour in a blocking solution containing 1X PBS and 5% normal goat serum. Slides were then incubated with 1:1000 mouse anti-Tyrosine Hydroxylase primary antibody (MAB318, Sigma-Aldrich, Darmstadt, Germany) in a PBS-T solution (PBS with 0.3% Triton) and 5% normal goat serum overnight, followed by a 2 hour incubation with 1:500 secondary antibody (Alexa Fluor 568, Invitrogen, Thermo Fisher Scientific, Waltham, MA). Slides were coverslipped with ProLong glass anti-fade mountant with NucBlue stain (Invitrogen, Thermo Fisher Scientific, Waltham, MA).

For each of the 11 brain regions identified above, immunostained slides were examined for the presence / absence of TH-positive cells at 20x magnification at 568nm fluorescent wavelength using a K5 camera (Leica Microsystems Inc., Deerfield, IL, USA). NucBlue (Hoechst 33342) stains nuclei, allowing for the identification of all 11 brain regions by comparing with our cresyl violet stained sections. We compared the presence / absence of TH-positive cell bodies in each of these regions to patterns of gene expression of TH across these brain regions in other teleosts ([19]; Table 1).

## Results

We developed an SDMN node atlas in stickleback, which includes 11 regions with important functions in social behavior across vertebrates [19]. Using serial, cresyl violet - stained sections, we identified cell groups based on the intensity of nuclear staining (Figure 1). We referenced the published brain atlases of zebrafish, guppy and cichlid to identify regions that correspond anatomically with the 11 teleost SDMN nodes [11,22–24] (Figure 2). To characterize nomenclature differences and highlight gaps in our understanding of the SDMN in teleosts, we then identified corresponding regions and sub-regions (or lack thereof) across atlases in an additional three fish species (Table 2). As our goal was to provide a neuroanatomical resource for stickleback with a particular focus on studying social behavior, we adopted the SDMN nomenclature used by [19]. Below we describe these anatomical regions and how they appear in stickleback, while highlighting similarities and differences to other teleosts. Next, we present results from TH immunohistochemistry to compare the presence of TH-positive cells in each region to other teleost lineages, demonstrating further variation among teleosts in this key gene product involved in dopamine synthesis. This variation underscores the importance of developing neuroanatomical resources in a variety of organisms, and highlights the diversity of brain morphology even within the teleost lineage.

**Fig. 1.**
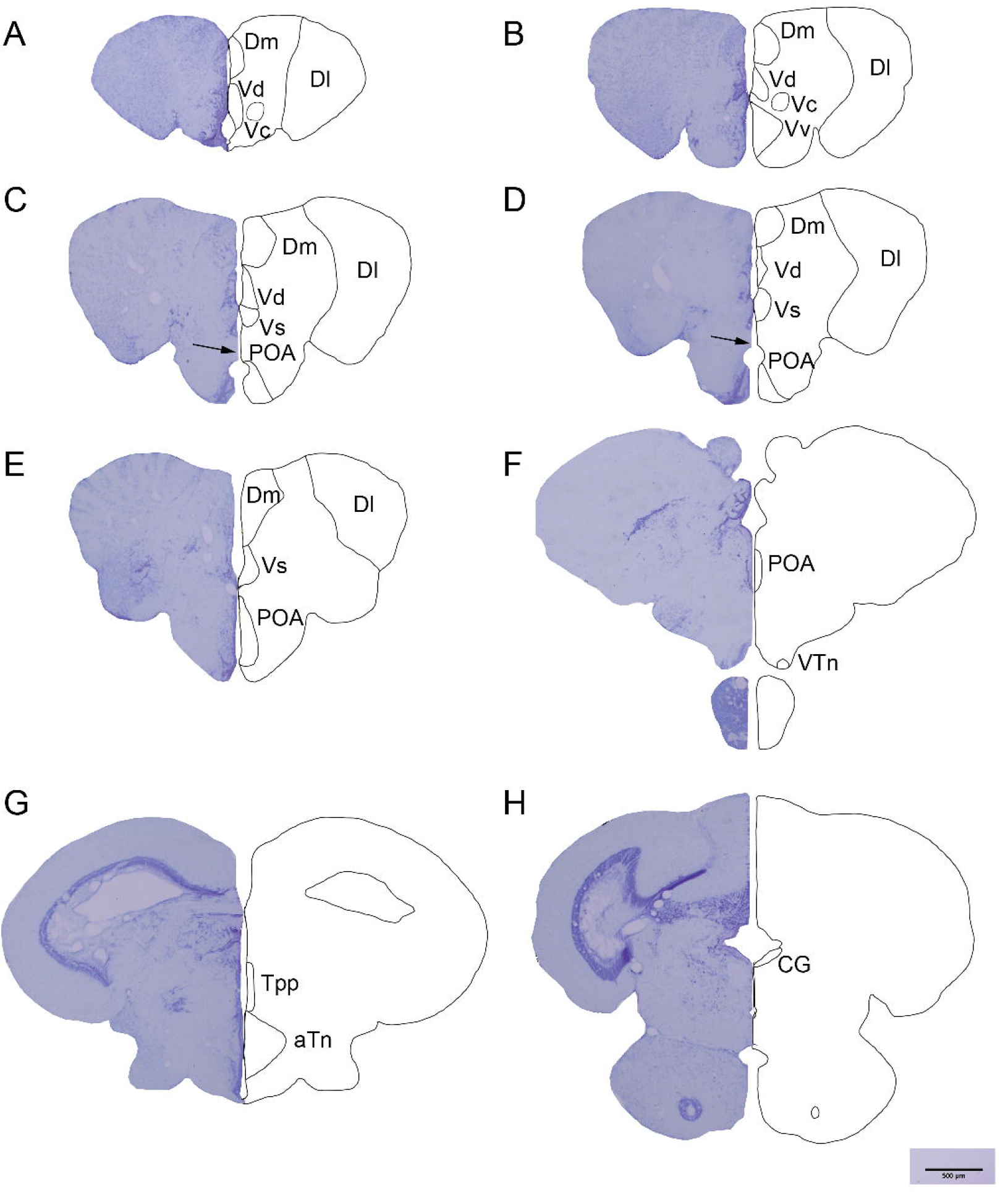
Transverse sections of the stickleback brain showing regions important for social behavior. Sections (A-H) proceed from rostral to caudal, with 500um scale bar shown at the bottom right. On the left of each panel is an image of the section stained with cresyl violet, and on the right is a line drawing outlining the location of each region. Medial zone of dorsal telencephalic area (Dm), lateral zone of dorsal telencephalic area (Dl), dorsal nucleus of V (Vd), central nucleus of V (Vc), ventral nucleus of V (Vv), supracommisural nucleus of V (Vs, V = ventral telencephalic area), preoptic area (POA), ventral tuberal nucleus (vTn), posterior tuberculum (Tpp), anterior tuberal nucleus (aTn), and central gray (CG). The arrow points to the anterior commissure, a landmark used to identify the start of the POA.

**Fig. 2.**
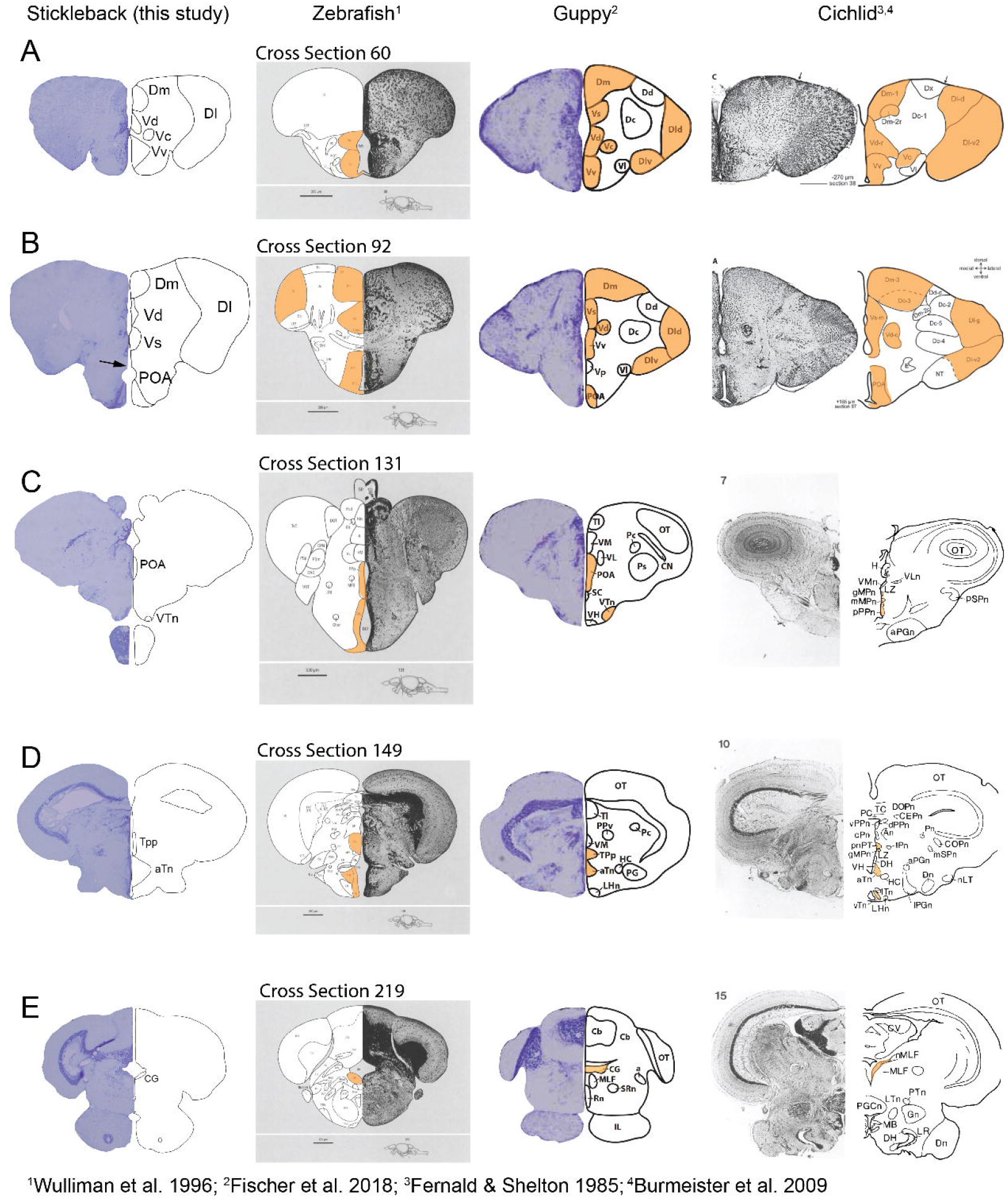
Comparison of brain regions important for social behavior delineated in stickleback (this study, left) to published brain atlases of three teleost species (zebrafish [11], guppy [22], and cichlid [23,24]), with corresponding regions in these other fishes colored in orange. Representative images (A-E) highlight similarities and differences in neuroanatomy and nomenclature across teleosts. The central gray (CG) in cichlid was not delineated in the published atlas shown here, but its location was inferred based on [82] and neuroanatomical similarities to other teleosts.

### Telencephalon

#### Dorsal telencephalic area (D)

The dorsal telencephalic area (D) is hypothesized to correspond to the pallium in other vertebrates, based on a model of developmental eversion [9,10]. As in other teleosts [11,22–24], this region consists of large masses of cells that make up the medial zone of D (Dm), the putative homolog of the mammalian basolateral amygdala, and the lateral zone of D (Dl), the putative homolog of the mammalian hippocampus (Figure 1A-E; Table 1). In other teleost brain atlases, these large regions are frequently subdivided (Table 1), but generally appear in similar anatomical positions across different atlases (Figure 2 A,B). In contrast to stickleback (this study), guppy and cichlid atlases, the zebrafish atlas shows the dorsal telencephalic area starting out more diffuse and organizes into discrete cell masses more caudally as the ventral telencephalic regions come in (Figure 2 A,B).

### Ventral telencephalic area (V)

The ventral telencephalic area (V), hypothesized to correspond to the subpallium in other vertebrates, is subdivided into the dorsal nucleus of V (Vd), central nucleus of V (Vc), ventral nucleus of V (Vv), and supracommissural nucleus of V (Vs) (Figure 1 A-E; Table 1). The Vd, the putative homolog of the mammalian nucleus accumbens, appears as a distinct cell group rostral to the anterior commissure, with the Vv, the putative homolog to the mammalian lateral septum, ventral to it and coming in as the Vd moves dorsally (Figure 1B). The Vc, the putative homolog to the mammalian striatum, appears as a small, circular region, coming in with the Vv (Figure 1 A,B). The start of the Vd, Vv, and Vc appears similar in other teleosts, however the Vs, the putative homolog of the mammalian bed nucleus of the stria terminalis / medial amygdala, comes in as the Vd moves dorsally in stickleback, but in guppy and cichlid Vs appears dorsal to the Vd, and appears serially after the Vd and Vv in zebrafish (Figure 2B).

### Diencephalon

#### Preoptic area (POA)

The preoptic area (POA) has been recently delineated in stickleback in a study showing its involvement in parental care behavior [34], and we maintain consistency with those delineations here. The start of the POA is easily identified anatomically, as it appears ventrally after a conspicuous gap of low cell density, the anterior commissure (Figure 1C), increases in size and moves dorsally (Figure 1D,E), and persists until a second gap of low cell density appears (Figure 1F). In other teleosts, the POA is frequently subdivided based on cell size, shape, and density (Figure 2C; Table 1).

#### Posterior tuberculum (Tpp)

The posterior tuberculum (Tpp), the putative homolog of the mammalian ventral tegmental area, comes in caudal to the POA, and corresponds with the optic tectum becoming larger and the tectal ventricle opening (Figure 1G). Anatomically, it appears similar across zebrafish, guppy, and cichlid, though there are variations in the shape and relative size (Figure 2D).

### Hypothalamus

The hypothalamus contains the anterior tuberal nucleus (aTn), the putative homolog of the mammalian ventromedial hypothalamus, and the ventral tuberal nucleus (vTn), the putative homolog of the anterior hypothalamus (Figure 1F,G; Table 1). While the aTn appears ventral to the Tpp and seems to be anatomically similar across teleosts (Figure 2D), the vTn was more difficult to find correspondence (Table 1). In part this may be due to nomenclature differences, with [19] referring to the ventral tuberal nucleus (vTn), while [32,33] refer to the ventral hypothalamus (Hv). Further confusing the matter, some atlases refer to the Hv or a corresponding anatomical region, the lateral tuberal nucleus (NLT), while others refer to the vTn (Table 1), and they appear in different anatomical positions, with the Hv and NLT closer to the midline, while the vTn appears more laterally (Figure 2C,D).

### Mesencephalon

#### Periaqueductal gray / central gray (CG)

The periaqueductal gray / central gray (CG) was identified as a distinct mass of cells bordering a ventricle as the optic tectum decreases in size and the cerebellum appears (Figure 1H). This appears consistent with its anatomical location in zebrafish and guppy, but was not delineated in the cichlid atlas we used (Figure 2E). A study characterizing the cichlid dopaminergic system showed a delineation of the periaqueductal gray (abbreviated PAG) more laterally [91], however the closest anatomical region to our atlas appears to be the nMLF (Figure 2E). The other three teleost atlases we examined did not delineate the CG / PAG, but we identified a potentially corresponding anatomical region in sea bass (Table 1).

#### Tyrosine hydroxylase (TH) quantification

We identified seven regions that had TH-positive cell bodies present: the Dm, Dl, Vd, Vc, POA, Tpp, and aTn (Figure 3; Table 1). When examining the distribution of TH immunoreactivity in other teleosts, there is quite a bit of variation in which regions show TH expression (Table 1) [19]. For example, while the Vc, POA, Tpp, and vTn generally show conserved expression in stickleback and the species surveyed in [19], TH expression in the aTn is typically absent in other teleosts, but was detected here (Figure 3F). Interestingly, TH expression in the Dm and Dl is typically absent in other teleosts and none of the 34 vertebrate species examined in [19] had the presence of dopamine-producing cell bodies in the Dm. Here, we did detect some cell body staining for TH in our study, albeit faint compared to staining in some other regions (Figure 3B,C). Additionally, the Vs, Vv, and CG typically show expression of TH in other teleosts, but we did not detect staining in these regions in our study (Table 1). Compared to other vertebrate lineages, the distribution of expression of TH appears particularly variable within teleosts [19], making this an interesting topic for future comparative study into functional variation in the SDMN with respect to social and motivated behaviors.

**Fig. 3.**
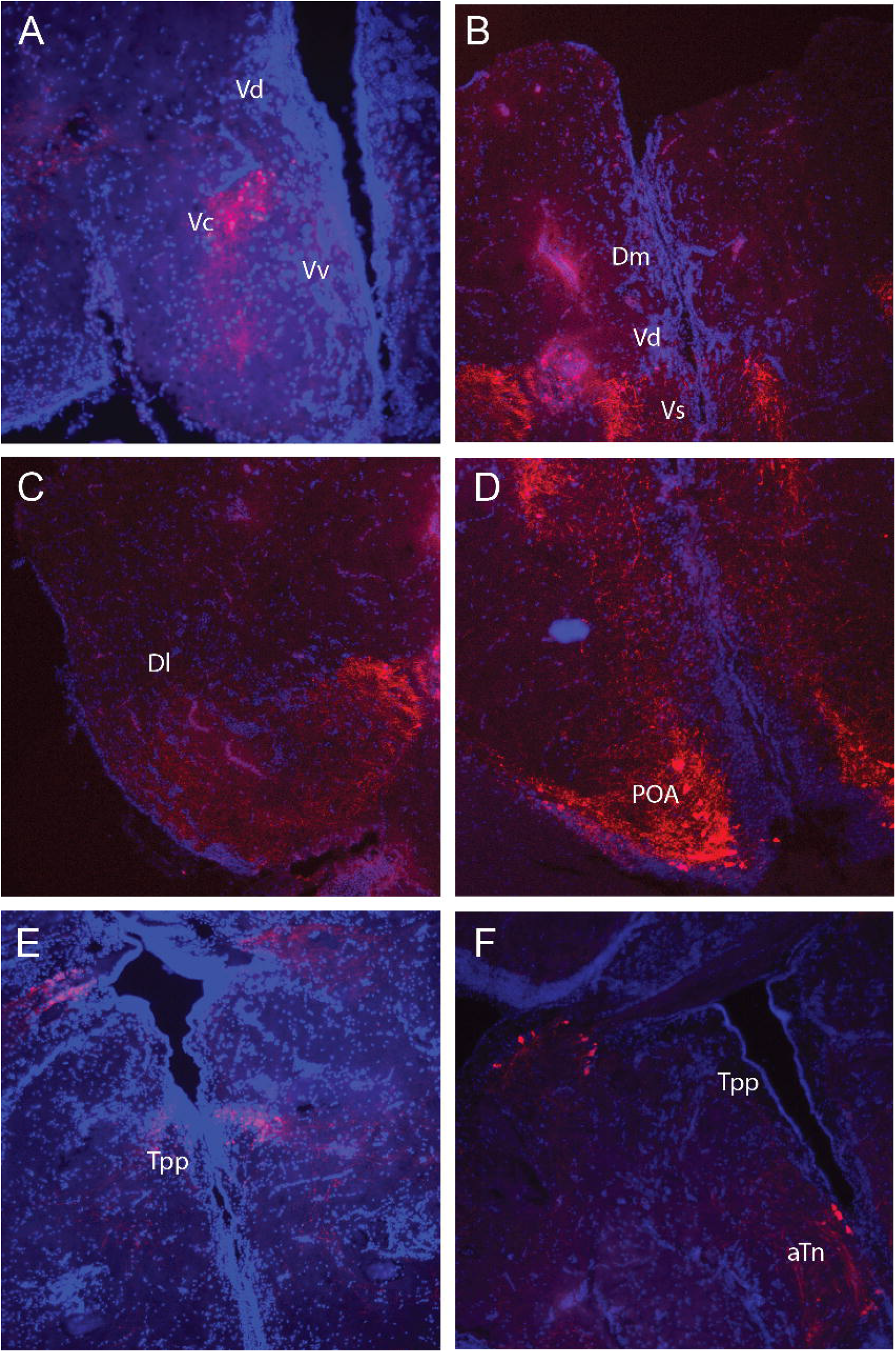
Images of select sections after tyrosine hydroxylase (TH) fluorescence immunohistochemistry. TH is the enzyme limiting dopamine synthesis and is a hallmark of hypothesized SDMN nodes [19]. Shown are images of the seven regions where the presence of TH-positive cell bodies was confirmed: medial zone of dorsal telencephalic area (Dm), lateral zone of D (Dl), central nucleus of the ventral telencephalic area (Vc), dorsal nucleus of V (Vd), preoptic area (POA), the periventricular nucleus of the posterior tuberculum (Tpp), and the anterior tuberal nucleus (aTn). Surrounding regions are labeled when present. Vd = dorsal nucleus of V, Vv = ventral nucleus of V. Fluorescent cell bodies are shown in red and nuclei stained with NucBlue are shown in blue.

## Discussion

In this study, we delineated in three-spined stickleback the 11 SDMN nodes that have been identified in teleosts. The anatomical locations of several regions were found to be similar across teleosts and easily identified in sticklebacks, most notably the POA, which has been shown to share functional similarities in other vertebrates [34]. However, we also identified some key differences, both anatomically and in the distribution of tyrosine hydroxylase immunoreactivity. The Vs and vTn were particularly difficult to identify due to variability in their locations across teleosts. The identification of the vTn was further confounded by the use of different nomenclature (Hv and NLT) across teleost atlases. Moreover, while the neuroanatomical locations of Dm and Dl in our atlas resemble those of other teleosts, the presence of cell body staining for tyrosine hydroxylase in these regions is typically absent in other teleosts but was detected in our study. Thus, neuroanatomical similarities across species may not necessarily reflect neurochemical or functional similarities, highlighting the need for molecular and functional characterizations of these regions to establish homology. Our SDMN atlas opens the way for future studies pertaining to the mechanisms of social behavior and social behavior evolution.

The neurobiological literature on sticklebacks to date has analyzed gross brain divisions such as the telencephalon and diencephalon (e.g. [35–38], or even whole brain (e.g. [39–41]). However, there is tremendous diversity in cell types and complex structure within the brain therefore future studies will benefit from resources that allow them to pinpoint specific brain areas and cell types important for social behavior. This SDMN atlas can facilitate future studies by providing a common reference for identifying brain regions, ensuring that brain area delineations and nomenclature are comparable across studies.

Our atlas’ utility is not limited to social behaviors, since the brain regions in the SDMN have been implicated in non-social behaviors as well. For example, the hippocampus and basolateral amygdala, putatively homologous to the Dl and the Dm in the teleost brain, respectively, are two areas in the SDMN but their function is not limited to social behavior. The hippocampus is involved in navigation and memory [42–46], and the teleost putative homolog the Dl has been similarly implicated [47–50]. Studies on stickleback have examined variation in brain size and morphology as a function of habitat complexity [51,52], and changes in Dl size have been hypothesized but not tested [52]. Such hypotheses could be tested with the help of a reference atlas. Relatedly, the amygdala is involved in fear responses and motivated behavior [53–57], and its putative teleost homolog the Dm is thought to serve a similar function [58–60]. Predator response, an ecologically relevant motivated behavior, is well-studied in stickleback [61–64], though few studies have examined the brain (but see [65,66]). Our SDMN atlas will enable better characterization of the Dm as a brain area mediating such responses.

Threespine sticklebacks exhibit tremendous diversity within and among populations in social behaviors such as courtship behavior and mate choice [67–69], territorial aggression and antipredator behavior [61–64,70–73] and parental care [74,75] and knowing the neurobiological causes and correlates of this behavioral variation has the potential to provide insights into the neural mechanisms underlying behavioral evolution. Marine sticklebacks are well-known for repeatedly moving into and rapidly adapting to new freshwater environments. The adaptive radiation of stickleback throughout the northern hemisphere provides numerous examples of convergent evolution of similar behavioral phenotypes in independent, replicated freshwater populations, where independently-derived populations share suites of morphological and behavioral adaptations that correlate with local environmental conditions [7]. Therefore the stickleback radiation provides opportunities to determine whether similar behaviors evolve via the same or different mechanisms and the extent to which neural circuit architecture promotes or constrains behavioral differences over evolutionary timescales [76]. Once there are more and better neuroanatomical resources available for them, sticklebacks are poised to be good subjects for understanding how genetic variation acts in or on neural circuits to influence behavioral variation [77].

An important consideration of this study is that different stickleback populations were used for identifying anatomically defined brain regions (an anadromous population in Japan) and for characterizing the presence/absence of TH staining (marine populations in Nova Scotia). The Japan population consists of the genetically divergent ‘Japan Sea’ and the ‘Pacific Ocean’ forms [25]. The individual used in this study was of the Pacific Ocean form, which diverged relatively recently (approximately 44 Kya) from the Atlantic stickleback, which includes Nova Scotia populations, and the Pacific and Atlantic Ocean forms are considered the same species [78]. We assume that our characterization of SDMN nodes is robust across populations because we do not expect there to be strong differences in brain organization among populations of the same species, and we found consistency in the neuroanatomical locations of SDMN nodes at least among the individuals and populations used in this study. Instead, we expect intraspecific brain differences to be more subtle, e.g., in connectivity, activity, and/or the relative size of specific brain areas important for different functions. Indeed, studies have described sex differences in brain size and morphology among stickleback populations [51,52,79,80] and the stickleback radiation provides many opportunities to learn about flexibility and constraint in brain structure at multiple levels of biological organization, from molecules to brain cells to circuits. A long term goal is to generate a pan-brain atlas that can act as a common reference frame for comparison of different patterns across individuals and populations; expanding the atlas to include data from different populations, sexes, and ages, will truly take advantage of the rich behavioral diversity of this system, as is being done in other natural systems such as cavefish [81].

In addition to a purely anatomical atlas, integration with resources such as brain cell type atlases and other molecular and neuroanatomical information will facilitate comparative and evolutionary studies. When the social behavior network (SBN), which is part of the larger SDMN, was hypothesized, brain regions were included in the network only when they met several criteria: the regions had to be reciprocally connected, express gonadal steroid receptors, and implicated in multiple social behaviors [20,21]. The evolution of the SDMN was evaluated via examination of gene products associated with dopaminergic system, sex steroid signaling, and nonapeptide systems [19]. In this manner, molecular information has informed cross-taxa comparisons and analyses; thus, the development of molecular atlases, built on top of neuroanatomical atlases, will help establish homologies and facilitate inter-taxa communication. Being able to perform comparative analysis across fish and other vertebrate lineages will enable researchers to link shared molecular pathways to a range of fundamental questions in behavior and brain evolution. Here, we demonstrated distinct differences in the distribution of tyrosine hydroxylase expression across SDMN nodes between sticklebacks and other fishes, highlighting the need for broader molecular characterizations to better understand the diversity of brain structure and function across species. The advent of species-agnostic sequencing technologies, such as single-cell/nuclei RNA sequencing, as well as spatial transcriptomic sequencing, has made it much more feasible to acquire molecular information for characterizing cell types in non-traditional model organisms. Integrated neuroanatomical resources will also enable the mapping of the entire connectome, or neural connections in the brain, to link brain structure to behavior and fitness.

The creation of these integrative molecular and neuroanatomical resources is an ambitious undertaking that will benefit from extensive collaboration. The phenotypic variation that is the stickleback system’s strength means that it will also take the global community of researchers, working on different stickleback populations and behaviors, to fully capture the diversity present in the brain of this system. In conclusion, while we presented a first step towards building a brain atlas in the form of a survey of the SDMN nodes, we hope that this will also be an invitation for researchers to contribute to the building of additional neuroanatomical resources.

## Statements

## Acknowledgements

Many thanks to E. Fischer for sharing her expertise and advice on generating a brain atlas. We also thank the Bell lab for helpful comments and suggestions on previous versions of this manuscript.

## Statement of Ethics

This study protocol was reviewed and approved by the Institutional Care and Use Committee at the University of Illinois Urbana Champaign, protocol number 24049.

## Author Contributions

TB: study concept and design, data acquisition, analysis and interpretation of data, drafting of the MS, revision of the MS, study supervision

UD: study concept and design, data acquisition, analysis and interpretation of data, drafting of the MS, revision of the MS, study supervision

CM: data acquisition, analysis and interpretation of data, analysis and interpretation of data, drafting of the MS, revision of the MS

JC: data acquisition, analysis and interpretation of data

MFM: study concept and design, data acquisition, analysis and interpretation of data, study supervision, revision of the MS

MK: data acquisition and interpretation, revision of the MS

NY: data acquisition and interpretation, revision of the MS

HK: data acquisition

JK: data acquisition

AMB: study concept and design, drafting of the MS, revision of the MS, administrative and material support, study supervision

## Data Availability Statement

All data used in this study are presented herein.

